# Raman imaging analysis of microvesicular and macrovesicular lipid droplets in non-alcoholic fatty liver disease

**DOI:** 10.1101/2020.07.20.212803

**Authors:** Takeo Minamikawa, Mayuko Ichimura-Shimizu, Hiroki Takanari, Yuki Morimoto, Ryosuke Shiomi, Hiroki Tanioka, Eiji Hase, Takeshi Yasui, Koichi Tsuneyama

## Abstract

Predominant evidence of non-alcoholic fatty liver disease (NAFLD) is the accumulation of excess lipids in the liver. A small group with NAFLD may have a more serious condition named non-alcoholic steatohepatitis (NASH). However, there is a lack of investigation of the accumulated lipids with spatial and molecular information. Raman microscopy has the potential to characterize molecular species and structures of lipids based on molecular vibration and can achieve high spatial resolution at the organelle level. We aim to demonstrate the feasibility of Raman microscopy for the investigation of NAFLD based on the molecular features of accumulated lipids. By applying the Raman microscopy to the liver of the NASH model mice, we succeeded in visualizing the distribution of lipid droplets (LDs) in hepatocytes. The detailed analysis of Raman spectra revealed the difference of molecular structural features of the LDs, such as the degree of saturation of lipids in the LDs. We also found that the inhomogeneous distribution of cholesterol in the LDs depending on the histology of lipid accumulation. We visualized and characterized the lipids of NASH model mice by Raman microscopy at organelle level. Our findings demonstrated that the Raman imaging analysis was feasible to characterize the NAFLD in terms of the molecular species and structures of lipids.

## Introduction

Non-alcoholic fatty liver disease (NAFLD) is the most common liver disease associated with the accumulation of excess lipids in the liver(1). The prevalence of NAFLD was increased worldwide and estimated to be 20 to 30% of the global population(2). NAFLD increases the risk of cirrhosis and liver cancer, and it is a disease with poor outcomes and limited therapeutic options in advanced stages. Furthermore, the clinical burden of NAFLD is not only in the liver dysfunctions but affects extrahepatic organs and regulatory pathways(3). Therefore, the development of diagnostic methods at an early stage and the understanding of the pathogenesis of NAFLD has become a serious clinical concern worldwide.

NAFLD can be classified into two types, *i.e*., non-alcoholic steatohepatitis (NASH) that leads to liver-related complications such as cirrhosis and hepatocellular carcinoma, and non-alcoholic fatty liver (NAFL) that basically has benign prognosis. Although several hypotheses were proposed to describe the pathogenesis of the NAFLD, the pathogenesis of the progression to NASH or NAFL is still under investigation. Moreover, NASH and NAFL at the early stage exhibit similar evidence of the accumulation of excess lipids without apparent inflammation and fibrosis, making diagnosis difficult. In the current studies, NAFLD was predominantly investigated in terms of protein or gene activities in the liver. Although the predominant evidence of NAFLD is the accumulation of excess lipids, the pathological role of the lipids on NAFLD has not been clearly understood due to a lack of lipid-based investigation of NAFLD.

To investigate NAFLD in terms of accumulated lipids, some visualization methods of lipid droplets (LDs) were proposed, *i.e*., dye-based visualization methods(4, 5), imaging mass spectrometry(6, 7) and Raman microscopy(8, 9). Among these methods, Raman microscopy is a good candidate for the investigation of the accumulated lipids based on molecular species and structures of lipids. Raman microscopy measures Raman spectra that reflect molecular vibrations of intrinsic molecules. Since the molecular vibrations are sensitive to the species of atoms and chemical bonds of molecules, Raman spectroscopy provides information about the molecular species and structures via molecular vibrations. As visible and tightly focused excitation laser can be used for observation, high spatial resolution down to 500 nm or smaller can be realized. Furthermore, it has the potential to utilize in vivo evaluation owing to the visible light usage. Raman microscopy is, therefore, a powerful tool for the investigation of lipids in cells and tissues(10–15). Although some researchers applied the Raman microscopy to NAFLD(16–18), the efficacy of Raman microscopy for the investigation of NAFLD is still under investigation.

In the present study, we seek to provide a proof-of-principle demonstration of Raman microscopy in the investigation of NAFLD. Especially, we visualize and characterize the accumulated lipids in the liver of mice by Raman microscopy for the investigation of NAFLD in terms of molecular species and structures of lipids.

## Materials and Methods

### NASH model animal

In this study, we utilized a fatty liver of mice exhibited NASH for the evaluation of Raman microscopy in the investigation of NAFLD. The detailed induction protocol and histopathological characterization of the NASH model mice used in this study were given in our previous study(19). NASH model mice were obtained by feeding six-weeks-old male TSOD mice (Institute for Animal Reproduction, Ibaraki, Japan) on a high-fat/cholesterol/cholate diet (iHFC diet #5; Hayashi Kasei, Osaka, Japan) and purified water for 26 weeks and housed under normal conditions. The liver of the NASH model mice was excised after euthanasia. All procedures performed in studies involving animals were in accordance with the ethical standards of the institution at which the studies were conducted, and ethical approval was obtained from the Institutional Animal Care and Use Committee of Tokushima University, Japan (Approval number: T30-32).

### Sample preparation

The excised liver was immediately embedded in 4% sodium carboxymethyl cellulose compound and snap-frozen in liquid nitrogen. The samples were stored at −80 °C until cryostat sectioning. The frozen samples were sliced into 5-μm thick sections with a cryostat microtome (Tissue Tek; Sakura, Tokyo, Japan). Two serial sections were obtained for Raman and histopathological analysis. The sections for the Raman analysis were mounted on a slide glass without any fixation nor staining. The sections for histopathological analysis were fixed with 95% ethanol and were subjected to hematoxylin and eosin (HE) staining. The histopathological characterization was performed by a pathologist (K.T.), who is the co-author of this article.

### Raman microscopy

Raman spectra and Raman spectral images were acquired with a home-built laser-scanning confocal Raman microscope with an imaging software (MwMapper; SicenceEdge Inc., Shizuoka, Japan). A single-mode frequency-doubled Nd:YAG laser (MSL-FN-532-S-100mW; CNI Laser, Changchun, China) operating at the wavelength of 532 nm was used as an excitation laser light. The excitation laser light was focused on a sample through a 10x objective lens (CFI Plan Apo Lambda 10X, 10x, NA = 0.45; Nikon, Tokyo, Japan) or a 60x objective lens (CFI Plan Apo Lambda 60XC, 60x, NA = 1.2; Nikon, Tokyo, Japan). The back-scattered Raman signal was collected with the same objective lens and detected by a spectrometer (IsoPlane 320, Princeton Instruments, Trenton, NJ, USA) with a cooled CCD image sensor (Pixis 400BR, −70°C, 1,340×400 pixels; Princeton Instruments, Trenton, NJ, USA). Two-dimensional Raman spectral images were obtained by scanning the laser focus. The excitation laser power and the exposure time were 10 mW on the sample plane and 0.1 s, respectively.

### Spectral preprocessing

Raman shifts of all Raman spectra were calibrated by using the known bands of a calibration lamp (IntelliCal; Princeton Instruments, Trenton, NJ, USA). To extract the Raman spectrum from a broad fluorescence background, we applied a modified polynomial curve fitting method(20). We estimated the autofluorescence component superposed on the Raman spectrum by calculating a modified least-squares 10-order polynomial curve with 100 iterations and then subtracted this polynomial from the raw spectra.

## Results

### Raman spectra of liver tissues of NASH model mice

Typical Raman spectra of liver tissues of NASH model mice were obtained to reveal the spectral features for Raman spectral analysis. We focused on macrovesicular LDs accumulated near a central vein, as shown in region A in Fig. 1a. The previous studies had shown that the crystallinity of LDs is an important indicator of NAFLD prognosis(21). By using the conventional polarization imaging with cross-Nicol configuration, the crystallinity of lipids in LDs can be identified as shown in Figs. 1b and 1c. We can observe a Maltese cross appearance owing to the polarization modulation by birefringence of a crystalline form of lipids, as indicated by the yellow arrowheads in Fig. 1c. If the Maltese cross is not apparent as indicated by the blue arrowhead in Fig. 1c, the LDs might predominantly contain an amorphous form of lipids. Thus, we investigate the spectral features of the Raman spectra of accumulated LDs with crystalline and amorphous forms of lipids in addition to hepatocytes with non-apparent LDs.

**Fig. 1.**
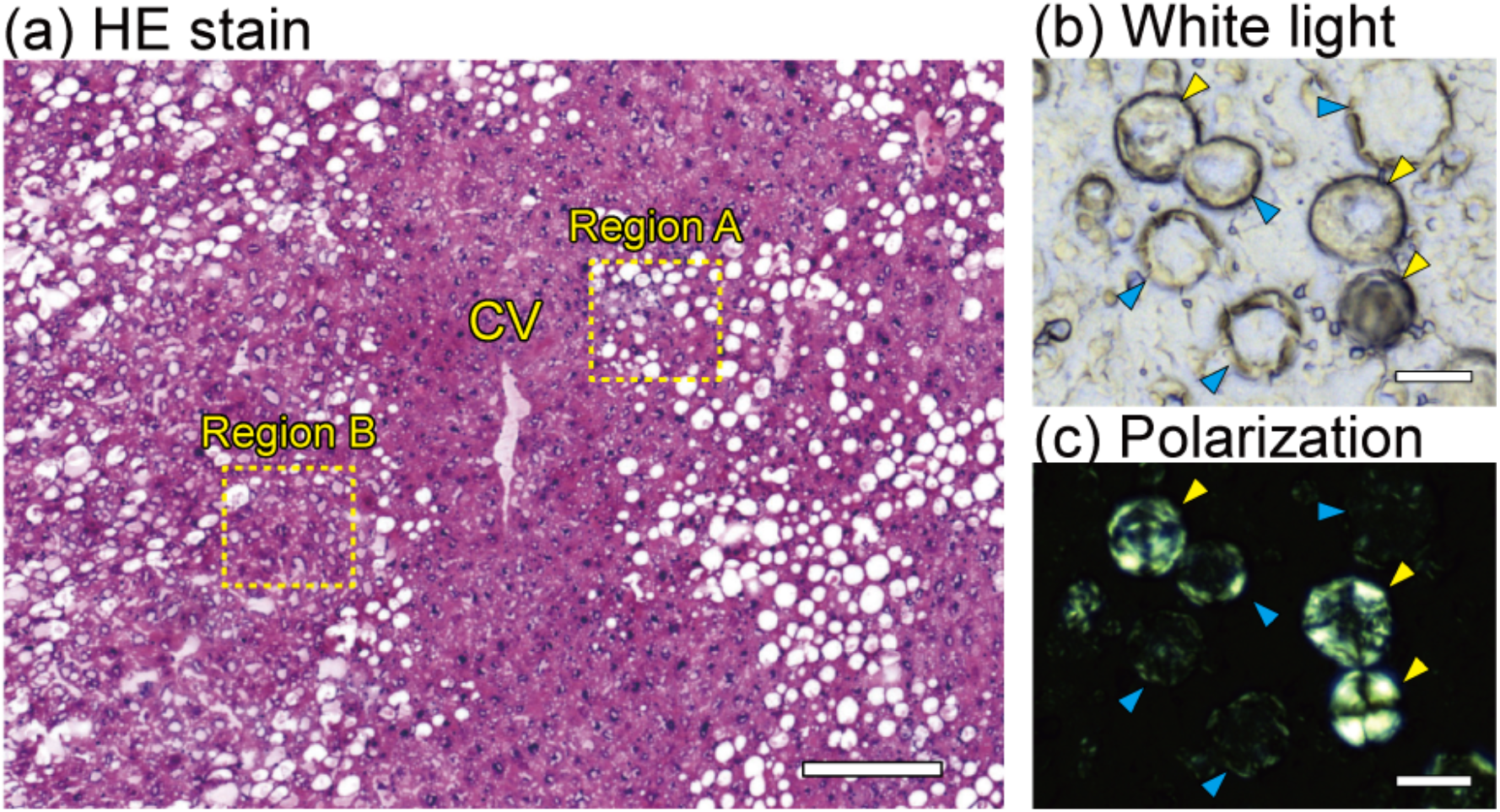
Typical histology of the liver tissue of a NASH model mouse. (a) HE-stained image of the NASH liver around a central vein (CV). (b) White light image of macrovesicular LDs of the NASH liver without any fixation nor staining. (c) Polarization image of the macrovesicular LDs of the NASH liver with cross-Nicol configuration. Blue arrowheads indicate LDs with an amorphous form of lipids. Yellow arrowheads indicate LDs with a crystalline form of lipids. Scale bars of (a), (b), and (c) indicate 200 μm, 10 μm, and 10 μm, respectively.

In the crystalline lipid-rich LDs, main Raman bands were found at 699, 1265, 1302, 1442, 1662, 1677, 1735, 2855, 2875, 2900, 2935, and 2964 cm^−1^ as shown in Fig. 2 and Table 1. The Raman bands at 1265, 1302, 1442, 2855, and 2900 cm^−1^ were assigned to CH2-related molecular bonds, indicating the presence of lipid contents in this region. The Raman band at 1662 cm^−1^ was assigned to C=C double bond stretching mode at the acyl chain, which is an indicator of the unsaturation of the acyl chain of lipids. The Raman bands at 699 and 1677 cm^−1^ respectively assigned to steroid ring vibration mode and C=C double bond stretching mode at carbon ring, indicating the presence of cholesterol content. As a result, the Raman spectra at the crystalline lipid-rich LDs reflected the molecular features of the LDs, including the cholesterols. The presence of cholesterol in the crystalline lipid-rich LDs was well agreed with the results of the fluorescence imaging with a cholesterol-specific dye in the previous studies(21).

**Fig. 2.**
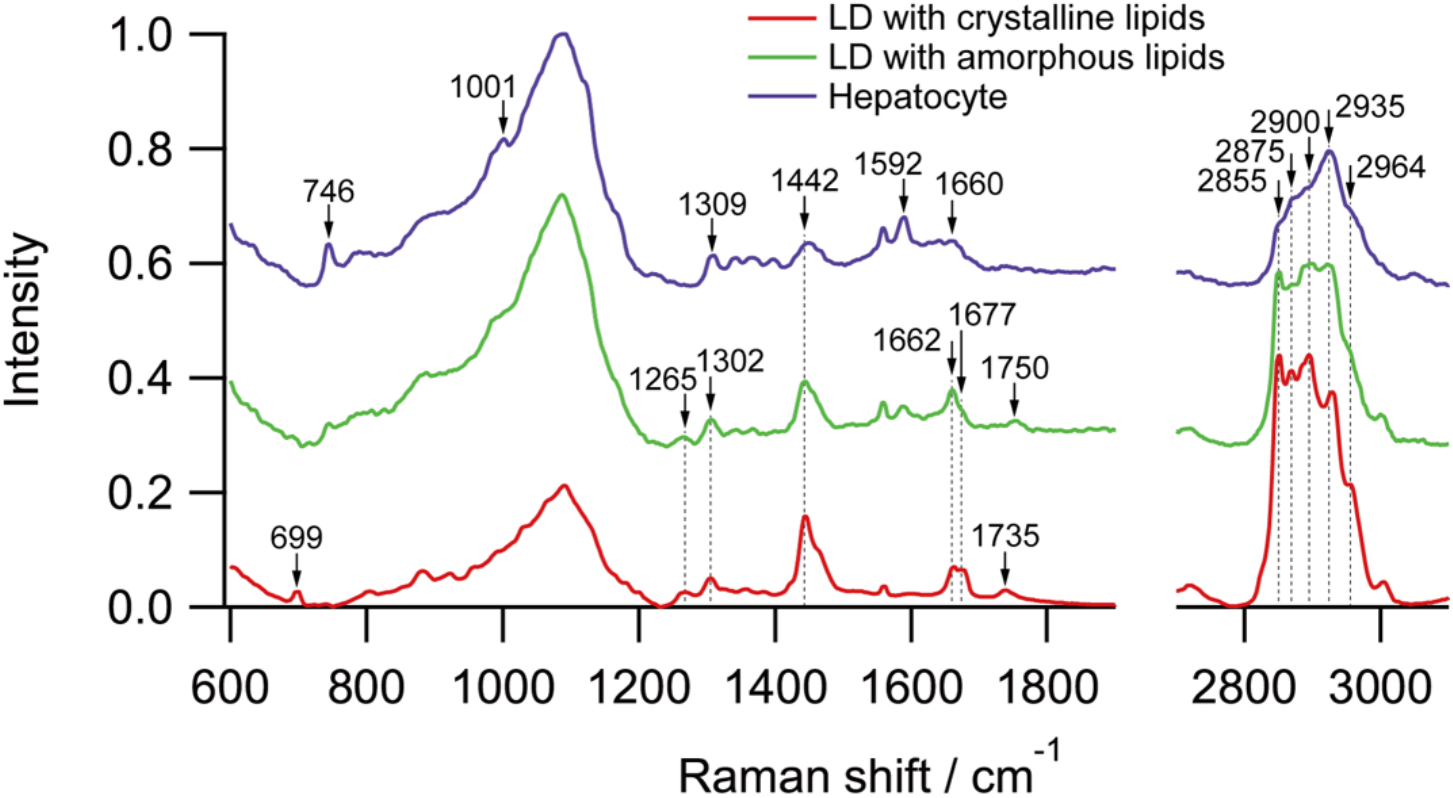
Typical Raman spectra of LDs with crystalline and amorphous forms of lipids and hepatocyte of the liver tissue of a NASH model mouse. The intensity was normalized by the highest intensities of the Raman spectra. LD, lipid droplet.

**Table 1.** Assignments of the Raman bands of LDs with crystalline and amorphous forms of lipids and hepatocyte of the liver tissue of a NASH model mouse.

| Assignment | LDs<br>(crystalline) | LDs<br>(amorphous) | Hepatocyte | Main<br>contribution |
| --- | --- | --- | --- | --- |
| Steroid rings stretch | 699 |  |  | Cholesterol |
| Pyrrole breathing |  |  | 746 | Heme |
| Si-O vibrations | 900-1200 | 900-1200 | 900-1200 | Background of<br>silica slide glass |
| Symmetric ring<br>breath<br>in phenylalanine |  |  | 1001 | Phenylalanine |
| =C-H bend | 1265 | 1265 |  | Lipids |
| CH<sub>2</sub> twist | 1302 | 1302 |  | Lipids |
| All heme bonds |  |  | 1309 | Heme |
| CH<sub>2</sub> bend | 1442 | 1442 | 1448 | Lipids |
| Redox and spin<br>state of heme |  |  | 1592 | Heme |
| Amide I |  |  | 1640-1680 | Proteins |
| C=C stretch<br>in acyl chain | 1662 | 1662 |  | Lipids |
| C=C stretch<br>in carbon ring | 1677 |  |  | Cholesterol |
| C=O stretch | 1735 | 1750 |  | Lipids |
| CH<sub>2</sub> symmetric<br>stretch | 2855 | 2855 | 2855 | Lipids |
| CH<sub>2</sub> asymmetric<br>stretch | 2875 | 2875 | 2875 | Lipids |
| Fermi resonance<br>CH<sub>2</sub> stretch | 2900 | 2900 | 2900 | Lipids |
| CH<sub>3</sub> symmetric<br>stretch | 2935 | 2935 | 2935 | Proteins |
| CH<sub>3</sub> asymmetric<br>stretch | 2964 | 2964 | 2964 | Proteins |

In the amorphous lipid-rich LDs, main Raman bands were found at 1265, 1302, 1442, 1662, 1750, 2855, 2875, 2900, 2935, and 2964 cm^−1^ as shown in Fig. 2 and Table 1. The main Raman bands were similar to that at the crystalline lipid-rich LDs, but the difference was found at 699 and 1677 cm^−1^, indicating relatively low cholesterol content at the LDs in terms of Raman spectral analysis. This result is also agreed with the result of the fluorescence imaging in the previous studies(21). Importantly, Raman microscopy additionally has the potential to quantitatively evaluate the relative content of molecular species and structures without staining by analyzing the relative intensity of each Raman band.

In contrast, in the hepatocytes with non-apparent LDs, main Raman bands were found at 746, 1001, 1309, 1448, 1592, 1660, 2855, 2875, 2900, 2935 and 2964 cm^−1^ as shown in Fig. 2 and Table 1. The Raman spectrum of the hepatocytes was different from that of these two types of LDs. Especially, heme-related Raman bands at 746, 1309, and 1592 cm^−1^ were observed in the hepatocytes, which might be derived by cytochromes present in mitochondria in hepatocytes or deposited hemoglobin. As a result, the Raman bands of LDs can be confirmed against that of hepatocytes, indicating the Raman spectral analysis of LDs can be performed in the liver tissues of the NASH model mice.

### Raman spectral imaging of macrovesicular LDs accumulated in hepatocytes

We investigated the spatial distribution of molecular contents of lipids in macrovesicular LDs accumulated in hepatocyte of liver tissues of NASH model mice. We obtained two-dimensional Raman images of macrovesicular LDs in region A in Fig. 1a. In this region, macrovesicular LDs with about 10 to 40 μm in diameter were diffusely distributed, as shown in Fig. 3a. The typical Raman spectra of LDs and hepatocytes in this region were shown in Fig. 3b. As similar results of Fig. 2, the Raman spectrum of LDs was predominantly composed of lipid-related Raman bands, such as 1442, 1662, 1677, and 2855 cm^−1^, as shown in Fig. 3b. Furthermore, cholesterol-related Raman bands such as 1677 cm^−1^ were also observed. Raman images of each typical Raman bands were shown in Fig. 3c. The lipid-related Raman bands, such as 1442, 1662, 1677, and 2855 cm^−1^, clearly reflect the distribution of LDs, which are well agreed with the histology of the liver tissues of the NASH model mice confirmed by the HE-stain image. As the imaging analysis, each LD seems to be separated from each other. In contrast, the Raman bands at 1592 and 2935 cm^−1^ visualized the distribution of hepatocytes or heme proteins.

To analyze the more specific molecular species and structures of LDs, we performed intensity ratio imaging by using lipid-related Raman bands. The intensity ratio calculation was performed at each pixel to obtain relative contributions of lipid species and structures with the normalization of the amount of lipids. Firstly, we obtained an intensity ratio of 1662 cm^−1^ against 2855 cm^−1^, as shown in Fig. 3d, which indicates the unsaturation degrees of lipids. The macrovesicular LDs were exhibited inhomogeneous distribution of the intensity ratio of 1662 cm^−1^ against 2855 cm^−1^. Notably, some particulate structures of the intensity ratio were found in the internal structure of LDs, as typically indicated by the arrowheads in Fig. 3d. These results might indicate the inhomogeneous distribution of lipids according to unsaturation degrees of lipids, even in the macrovesicular LDs with morphologically uniform distribution.

**Fig. 3.**
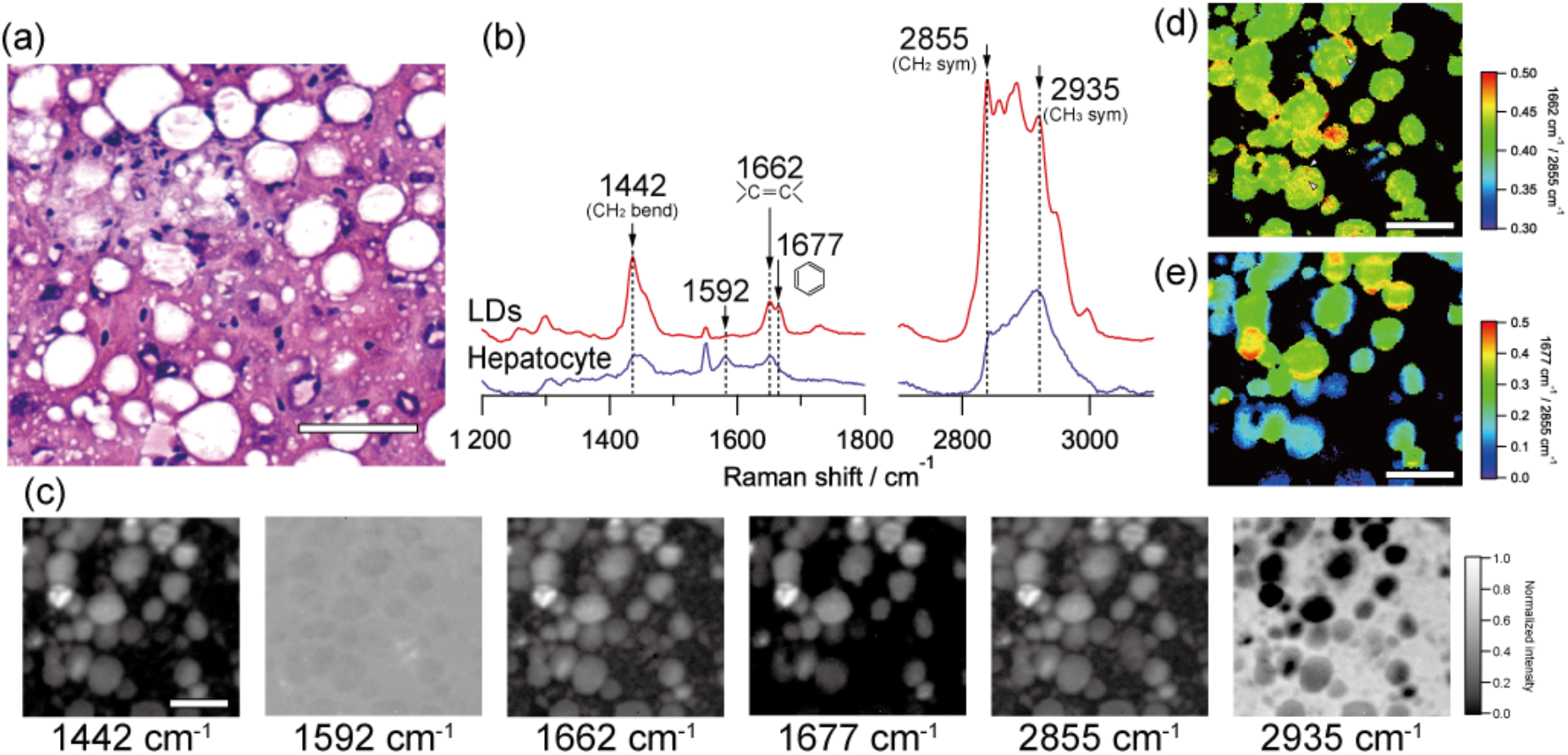
Raman spectral imaging of macrovesicular LDs accumulated in hepatocyte of liver tissues of NASH model mice. (a) HE-stained image of macrovesicular LDs indicated by region A in Fig. 1a. Raman imaging was performed on an unstained tissue section corresponding to the same position as the HE-stained image. (b) Typical Raman spectra of LDs and hepatocytes in this region. (c) Raman images of typical Raman bands. (d) Intensity ratio imaging of Raman bands of 1662 and 2855 cm^−1^, indicating the unsaturation degree of LDs. (e) Intensity ratio imaging of Raman bands of 1677 and 2855 cm^−1^, indicating the relative content of cholesterol. These intensity ratio images were obtained at the region with enough high intensity at the Raman band of 2855 cm^−1^, while the other region was masked. Scale bars of (a), (c), (d), and (e) indicate 50 μm.

We also obtained the intensity ratio of 1677 cm^−1^ against 2855 cm^−1^, as shown in Fig. 3e, which indicates the relative content of cholesterol. We found that the inhomogeneous distribution of the intensity ratio of 1677 cm^−1^ against 2855 cm^−1^ among LDs. In addition, some of the LDs exhibited a high-intensity ratio partially on the outer wall of the LDs. These results might indicate that the relative content of cholesterol was varied depending on the macrovesicular LDs, and the cholesterol distribution was unevenly distributed even in a single macrovesicular LD.

### Raman spectral imaging of microvesicular LDs accumulated in hepatocytes

We also investigated the spatial distribution of molecular contents of lipids in microvesicular LDs accumulated in hepatocyte of liver tissues of NASH model mice. We obtained two-dimensional Raman images of macrovesicular LDs at the dashed squared region of Fig. 4a that are in region B in Fig. 1a. In this region, microvesicular LDs with about a few micrometers or less in diameter were diffusely distributed, as shown in Fig. 4a. The typical Raman spectra of LDs and hepatocytes in this region were shown in Fig. 4b. The Raman spectrum of LDs was exhibited similar to the macrovesicular LDs, such as 1442, 1662, 1677, and 2855 cm^−1^, as shown in Fig. 3b. However, the relative contribution of 1677 cm^−1^ was much smaller than that of macrovesicular LDs. Raman images of each typical Raman bands were shown in Fig. 4c. The lipid-related Raman bands, such as 1442, 1662, 1677, and 2855 cm^−1^, reflect the distribution of LDs, but interestingly, we found the connecting structure between LDs, which was not apparent in the HE-stained image. In contrast, the Raman bands at 1592 and 2935 cm^−1^ visualized the distribution of hepatocytes or heme proteins as similar to the result around the macrovesicular LDs.

**Fig. 4.**
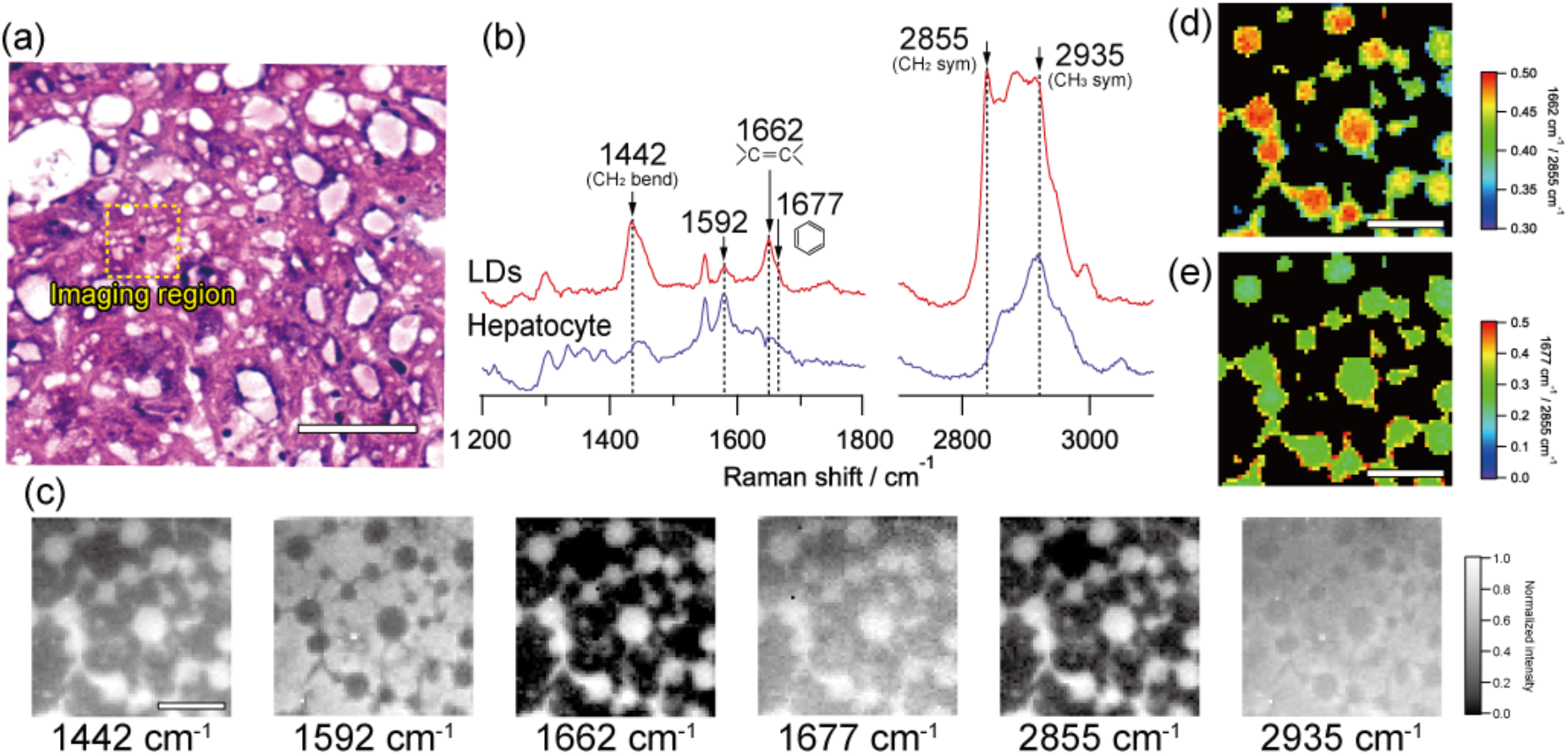
Raman spectral imaging of microvesicular LDs accumulated in hepatocyte of liver tissues of NASH model mice. (a) HE-stained image of microvesicular LDs indicated by region B in Fig. 1a. Raman imaging was performed on an unstained tissue section corresponding to the dashed squared region of the HE-stained image. (b) Typical Raman spectra of LDs and hepatocytes in this region. (c) Raman images of typical Raman bands. (d) Intensity ratio imaging of Raman bands of 1662 and 2855 cm^−1^, indicating the unsaturation degree of LDs. (e) Intensity ratio imaging of Raman bands of 1677 and 2855 cm^−1^, indicating the relative content of cholesterol. These intensity ratio images were obtained at the region with enough high intensity at the Raman band of 2855 cm^−1^, while the other region was masked. Scale bars of (a), (c), (d), and (e) indicate 50 μm, 10 μm, 10 μm, and 10 μm, respectively.

Intensity ratio imaging was also performed by using lipid-related Raman bands. We obtained the intensity ratio of 1662 cm^−1^ against 2855 cm^−1^, as shown in Fig. 3d, which indicates the unsaturation degrees of lipids. The microvesicular LDs have exhibited an almost homogeneous distribution of the intensity ratio of 1662 cm^−1^ against 2855 cm^−1^. We also obtained the intensity ratio of 1677 cm^−1^ against 2855 cm^−1^, as shown in Fig. 4e, which indicates the relative content of cholesterol. The distribution of the intensity ratio of 1677 cm^−1^ against 2855 cm^−1^ was also homogeneous. These results might indicate the unsaturation degree of microvesicular LDs, and the relative content of cholesterol could distribute homogeneously in microvesicular LDs.

### Statistical comparison of molecular species and structural features of LDs between microvesicular and macrovesicular LDs

Finally, we investigated the difference of molecular species and structural features of LDs between microvesicular and macrovesicular LDs. We evaluated the intensity ratio of 1662 cm^−1^ against 2855 cm^−1^ and the that of 1677 cm^−1^ against 2855 cm^−1^ of the central region of the 37 LDs in macrovesicular LDs and 21 LDs in microvesicular LDs. The intensity ratio analyses of these Raman bands were shown in Fig. 5. The mean intensity ratio of 1662 cm^−1^ against 2855 cm^−1^ of the microvesicular LDs (0.438 ± 0.038) was significantly higher than that of the macrovesicular LDs (0.425 ± 0.034). This result shows the microvesicular LDs have a higher unsaturation degree of lipids than the macrovesicular LDs. The mean intensity ratio of 1677 cm^−1^ against 2855 cm^−1^ of the microvesicular LDs (0.308 ± 0.061) was also significantly higher than that of the macrovesicular LDs (0.218 ± 0.104). This result shows the microvesicular LDs have a higher cholesterol content on average. However, the highest intensity ratio of 1677 cm^−1^ against 2855 cm^−1^ was found in the macrovesicular LDs, showing the partial aggregation of a higher content of cholesterol.

**Fig. 5.**
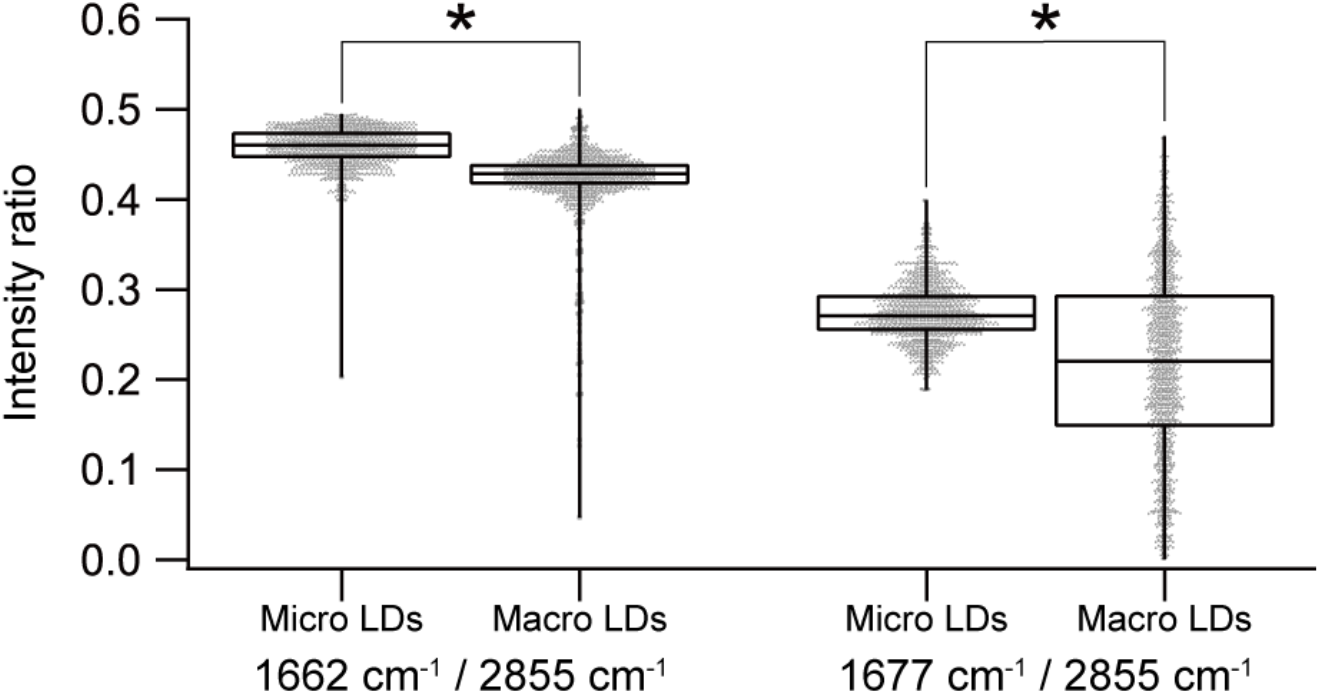
Box plot analysis of intensity ratio of Raman bands indicating the unsaturation degree of lipids (1662 cm^−1^/2855 cm^−1^) and the relative content of cholesterol (1677 cm^−1^/2855 cm^−1^) of microvesicular LDs (Micro LDs) and macrovesicular LDs (Macro LDs). A total of 1270 points for each of 21 microvesicular LDs and 37 macrovesicular LDs were evaluated. The individual data are shown by dots. *, significant difference (p < 0.01).

## Discussion

In the present study, we evaluated liver tissues of NASH model mice in terms of the molecular analysis of LDs within hepatocytes. We analyzed spectral features of the Raman spectra of LDs with crystalline and amorphous forms of lipids and hepatocytes. We also successfully visualized the LDs according to the molecular species and structural features of lipids in the microvesicular and macrovesicular LDs. The Raman microscopy reveals the complicated distributions of lipids in LDs, which cannot be identified by conventional histological imaging techniques. Our results provided a proof-of-principle for the investigation of NAFLD using Raman microscopy in terms of the molecular species and structures of accumulated lipids in NAFLD.

NAFLD induces hepatocyte steatosis and liver dysfunction with glycolipid metabolic disorders based on insulin resistance. The pathogenesis is thought to be due to various factors such as oxidative stress, endoplasmic reticulum stress, autophagy, lipotoxicity owing to free fatty acids, and innate immune activation associated with bacterial intestinal abnormalities(22–24); accordingly, various researches on NAFLD were conducted in terms of genome-wide association studies(25), molecular biological studies(23, 26, 27), and medical studies in both the aspect of treatment and diagnosis(28, 29). As regarding lipids, abnormal lipid metabolism and the induction of inflammation by overproduction of free fatty acids have been widely investigated. However, the current studies related to lipids have been predominantly performed by ensemble-averaged analysis of lipids without spatial information or by morphological analysis of accumulated LDs without molecular information. Since the accumulation of excess lipids is fundamental evidence of NAFLD, the analysis of lipids with both spatial and molecular information must provide an essential perspective in elucidating the pathogenesis of NAFLD. Our results demonstrated the feasibility of Raman microscopy for the molecular analysis with spatial information, and the Raman microscopy may provide an effective means to investigate the pathogenesis of NAFLD based on the characteristics of accumulated lipids.

Conventional bioimaging methods focusing on lipids are dye-based visualization methods and imaging mass spectrometry; however, the molecular selectivity of the dye-based visualization methods in lipid analysis is low because of the difficulty of the application of such as an immune-staining method or similar method of fluorescent proteins in the same manner of protein imaging. Therefore, the dye-based visualization of lipids for NAFLD analysis is limited to morphological analysis. The imaging mass spectrometry is sensitive and selective in terms of lipid molecules and can clarify the details of lipid species, the length of acyl chains, and the degree of unsaturation. However, it is challenging to analyze microvesicular LDs or internal structures of LDs due to its low spatial resolution (> 10 μm). By contrast, Raman microscopy can achieve very high spatial resolution up to about half of the excitation wavelength, while obtaining molecular information via molecular vibrations. Moreover, Raman microscopy has a potential for application in intraoperative use by using a surgical microscope or flexible optical fiber system(10, 30). Raman microscopy offers the advantages of fast, in situ, non-destructive and molecular vibration-based observation, enabling the diagnosis applications for NAFLD without any treatment such as fixation nor staining. Although further development of an intraoperative Raman microscopy system is required, our proposed approach may also provide a unique and powerful means for NAFLD diagnosis in the future.

## Conclusion

In conclusion, we demonstrated Raman spectral imaging of the liver tissues of NASH model mice. Our results revealed the feasibility of the Raman microscopy for the analysis of NAFLD based on the molecular features of accumulated LDs. Although further studies are required, our proposed approach may provide new insights into the investigation of the pathogenesis of NAFLD and the new development for NAFLD diagnosis based on accumulated lipids in the future.

## Data availability statement

The data that support the findings of this study are available from the corresponding author upon reasonable request.

## Acknowledgement

We acknowledge Ms. Natsuko Takeichi and Ms. Asaka Murakami of Tokushima University for their help in English proofreading of the manuscript. This work was supported by PRESTO program (JPMJPR17PC) of Japan Science and Technology Agency (JST), Japan, a Grant-in-Aid for Challenging Research (Exploratory) (JP19K22969) from the Japan Society for the Promotion of Science (JSPS), Japan, a research grant from Kowa Life Science Foundation, Japan, a research grant from the Research Clusters program of Tokushima University (1802003), Japan.

## Abbreviations

HE: Hematoxylin and eosin
LD: Lipid droplet
Macro LD: Macrovesicular lipid droplet
Micro LD: Microvesicular lipid droplet
NAFL: Non-alcoholic fatty liver
NAFLD: Non-alcoholic fatty liver disease
NASH: Non-alcoholic steatohepatitis

## Reference

1. Angulo, P. 2002. Nonalcoholic fatty liver disease. N. Engl. J. Med. 346: 1221–1231.

2. Younossi, Z. M., A. B. Koenig, D. Abdelatif, Y. Fazel, L. Henry, and M. Wymer. 2016. Global epidemiology of nonalcoholic fatty liver disease-Meta-analytic assessment of prevalence, incidence, and outcomes. Hepatology 64: 73–84.

3. Byrne, C. D., and G. Targher. 2015. NAFLD: a multisystem disease. J. Hepatol. 62: S47–64.

4. Gocze, P. M., and D. A. Freeman. 1994. Factors underlying the variability of lipid droplet fluorescence in MA-10 Leydig tumor cells. Cytometry 17: 151–158.

5. Spandl, J., D. J. White, J. Peychl, and C. Thiele. 2009. Live cell multicolor imaging of lipid droplets with a new dye, LD540. Traffic 10: 1579–1584.

6. Debois, D., M. P. Bralet, F. Le Naour, A. Brunelle, and O. Laprevote. 2009. In situ lipidomic analysis of nonalcoholic fatty liver by cluster TOF-SIMS imaging. Anal. Chem. 81: 2823–2831.

7. Alamri, H., N. H. Patterson, E. Yang, P. Zoroquiain, A. Lazaris, P. Chaurand, and P. Metrakos. 2019. Mapping the triglyceride distribution in NAFLD human liver by MALDI imaging mass spectrometry reveals molecular differences in micro and macro steatosis. Anal Bioanal Chem 411: 885–894.

8. van Manen, H. J., Y. M. Kraan, D. Roos, and C. Otto. 2005. Single-cell Raman and fluorescence microscopy reveal the association of lipid bodies with phagosomes in leukocytes. Proc. Natl. Acad. Sci. U. S. A. 102: 10159–10164.

9. Slipchenko, M. N., T. T. Le, H. Chen, and J. X. Cheng. 2009. High-speed vibrational imaging and spectral analysis of lipid bodies by compound Raman microscopy. J Phys Chem B 113: 7681–7686.

10. Hattori, Y., Y. Komachi, T. Asakura, T. Shimosegawa, G. Kanai, H. Tashiro, and H. Sato. 2007. In vivo Raman study of the living rat esophagus and stomach using a micro-Raman probe under an endoscope. Appl. Spectrosc. 61: 579–584.

11. Giarola, M., B. Rossi, E. Mosconi, M. Fontanella, P. Marzola, I. Scambi, A. Sbarbati, and G. Mariotto. 2011. Fast and minimally invasive determination of the unsaturation index of white fat depots by micro-Raman spectroscopy. Lipids 46: 659–667.

12. Okada, M., N. I. Smith, A. F. Palonpon, H. Endo, S. Kawata, M. Sodeoka, and K. Fujita. 2012. Label-free Raman observation of cytochrome c dynamics during apoptosis. Proc. Natl. Acad. Sci. U. S. A. 109: 28–32.

13. Minamikawa, T., Y. Harada, and T. Takamatsu. 2015. Ex vivo peripheral nerve detection of rats by spontaneous Raman spectroscopy. Sci. Rep. 5: 17165.

14. Stiebing, C., L. Schmolz, M. Wallert, C. Matthaus, S. Lorkowski, and J. Popp. 2017. Raman imaging of macrophages incubated with triglyceride-enriched oxLDL visualizes translocation of lipids between endocytic vesicles and lipid droplets. J. Lipid Res. 58: 876–883.

15. Yamamoto, T., T. Minamikawa, Y. Harada, Y. Yamaoka, H. Tanaka, H. Yaku, and T. Takamatsu. 2018. Label-free evaluation of myocardial infarct in surgically excised ventricular myocardium by Raman spectroscopy. Sci. Rep. 8: 14671.

16. Le, T. T., A. Ziemba, Y. Urasaki, S. Brotman, and G. Pizzorno. 2012. Label-free evaluation of hepatic microvesicular steatosis with multimodal coherent anti-Stokes Raman scattering microscopy. PLoS One 7: e51092.

17. Kochan, K., E. Maslak, C. Krafft, R. Kostogrys, S. Chlopicki, and M. Baranska. 2015. Raman spectroscopy analysis of lipid droplets content, distribution and saturation level in Non-Alcoholic Fatty Liver Disease in mice. J Biophotonics 8: 597–609.

18. Helal, K. M., J. N. Taylor, H. Cahyadi, A. Okajima, K. Tabata, Y. Itoh, H. Tanaka, K. Fujita, Y. Harada, and T. Komatsuzaki. 2019. Raman spectroscopic histology using machine learning for nonalcoholic fatty liver disease. FEBS Lett. 593: 2535–2544.

19. Ichimura, M., M. Kawase, M. Masuzumi, M. Sakaki, Y. Nagata, K. Tanaka, K. Suruga, S. Tamaru, S. Kato, K. Tsuneyama, and K. Omagari. 2015. High-fat and high-cholesterol diet rapidly induces non-alcoholic steatohepatitis with advanced fibrosis in Sprague-Dawley rats. Hepatol. Res. 45: 458–469.

20. Lieber, C. A., and A. Mahadevan-Jansen. 2003. Automated Method for Subtraction of Fluorescence from Biological Raman Spectra. Appl. Spectrosc. 57: 1363–1367.

21. Ioannou, G. N., W. G. Haigh, D. Thorning, and C. Savard. 2013. Hepatic cholesterol crystals and crown-like structures distinguish NASH from simple steatosis. J. Lipid Res. 54: 1326–1334.

22. Marra, F., and S. Lotersztajn. 2013. Pathophysiology of NASH: perspectives for a targeted treatment. Curr. Pharm. Des. 19: 5250–5269.

23. Arab, J. P., M. Arrese, and M. Trauner. 2018. Recent Insights into the Pathogenesis of Nonalcoholic Fatty Liver Disease. Annu. Rev. Pathol. 13: 321–350.

24. Buzzetti, E., M. Pinzani, and E. A. Tsochatzis. 2016. The multiple-hit pathogenesis of nonalcoholic fatty liver disease (NAFLD). Metabolism 65: 1038–1048.

25. Yu, J., S. Marsh, J. Hu, W. Feng, and C. Wu. 2016. The Pathogenesis of Nonalcoholic Fatty Liver Disease: Interplay between Diet, Gut Microbiota, and Genetic Background. Gastroenterol. Res. Pract. 2016: 2862173.

26. Berlanga, A., E. Guiu-Jurado, J. A. Porras, and T. Auguet. 2014. Molecular pathways in nonalcoholic fatty liver disease. Clin. Exp. Gastroenterol. 7: 221–239.

27. Malaguarnera, M., M. Di Rosa, F. Nicoletti, and L. Malaguarnera. 2009. Molecular mechanisms involved in NAFLD progression. J. Mol. Med. (Berl.) 87: 679–695.

28. Romero-Gomez, M., S. Zelber-Sagi, and M. Trenell. 2017. Treatment of NAFLD with diet, physical activity and exercise. J. Hepatol. 67: 829–846.

29. Ali, R., and K. Cusi. 2009. New diagnostic and treatment approaches in non-alcoholic fatty liver disease (NAFLD). Ann. Med. 41: 265–278.

30. Buschman, H. P., E. T. Marple, M. L. Wach, B. Bennett, T. C. Schut, H. A. Bruining, A. V. Bruschke, A. van der Laarse, and G. J. Puppels. 2000. In vivo determination of the molecular composition of artery wall by intravascular Raman spectroscopy. Anal. Chem. 72: 3771–3775.

